# Limitations of line-scan MRI for directly measuring neural activity

**DOI:** 10.1101/2023.11.03.565394

**Authors:** Joshua M. Wilson, Hua Wu, Adam B. Kerr, Brian A. Wandell, Justin L. Gardner

## Abstract

Several groups have reported using 2D line-scan MRI sequences in humans and mice to directly measure neural responses to stimuli (the “DIANA response”). Other groups have been unable to replicate the DIANA response, even with higher field strength and more repetitions. Part of this discrepancy is likely due to a limited understanding of the noise profile of the line-scan MRI sequence: specifically, the consequences of deviations from the assumption of stationarity between each line acquisition. Here, we collected data using an MRI line-scan method while human subjects viewed a blank screen with the purpose of studying noise unique to the acquisition sequence. We found temporal fluctuations in the reconstructed time series from localized groups of voxels that could easily be confused with neural responses to stimuli. These fluctuations were present both in the head and in the surrounding empty volume along the span of the phase-encoding direction from the head. The timing of these fluctuations varied systematically and smoothly along the phase-encoding direction. These features can be explained by a model that accounts for the acquisition sequence and incorporates time-varying contrast fluctuations in the imaging substrate. Using the model, we quantify the amount of cortical- and scan-averaging one might need to reliably distinguish a DIANA response from noise.

## Introduction

There has been a decades-long search for a noninvasive magnetic resonance method that directly measures neural responses at high spatio-temporal resolution (Bandettini et al., 2005; Bodurka and Bandettini, 2002; Petridou et al., 2006). Recent studies reported that line-scan MRI imaging sequences (Bohning et al., 1990; Lee et al., 2021; Silva and Koretsky, 2002; Waterton et al., 1985) can record thalamic and cortical electrical responses to stimuli in mice (Toi et al., 2022) and humans (Zhang et al., 2023). The recorded MRI time series appear to match electrophysiological measurements at millisecond resolution. However, other groups following similar protocols have been unable to replicate these results in both mice and humans (Choi et al., 2023; Hodono et al., 2023; Thorp, 2023).

One potential reason for this discrepancy among groups is that the signal sizes are small compared to other functional MRI measurements, such as BOLD (Ogawa et al., 1992, 1990). Equally important, the noise characteristics of the acquisition sequence have not been well-characterized, which makes it difficult to confidently discriminate signal from noise. Lack of information about the noise limits our ability to set measurement parameters such as the number of repeats and the size of spatial averaging region required to achieve high signal-to-noise ratios (SNR).

The long acquisition sequence used to reconstruct images with the line-scanning technique is particularly susceptible to artifacts due to violations of stationarity during the acquisition. The line-scanning technique acquires each line of k-space repeatedly during and after stimulus presentation, for a duration (line duration) typically on the order of a second as required to capture a physiological response. This allows for fast temporal resolution, because each single line of k-space is acquired rapidly, typically with a repetition time (TR) of a few milliseconds. Images are reconstructed by combining all the lines of k-space of each image, captured after different stimulus presentations. The fundamental assumption is stationarity; across different stimulus presentations the brain tissue is assumed to be in the same state, thus allowing for combination of different lines of k-space into images even though they were collected at different times. This approach contrasts with conventional Echo-Planar Imaging (EPI) imaging (Mansfield, 1977; Stehling et al., 1991), in which all k-space lines of a single image are collected before moving to the next image. This line-scan approach increases the acquisition time of any individual image by orders of magnitude; depending on the line duration, number of lines of k-space, TR, and other scan parameters, one can expect that any individual image is collected over a period of 10-60 seconds. This long acquisition time greatly increases the chance that subject head motion, respiration and cardiac signals (GOTO et al., 2016; Hermes et al., 2023; Hu et al., 1995; Liu, 2016; Network et al., 2013), as well as fluctuations in brain state not linked to stimulus presentation (Biswal et al., 1995; Chang et al., 2016; Lurie et al., 2020) can violate the assumption of stationarity across different stimulus presentations. These nonstationarities between line acquisitions can cause artifacts that will not be restricted to specific locations; because each line of k-space contributes to all points in the image, nonstationarities between collections of different k-space lines can result in artifacts spread across the whole image.

Here we quantify the unique spatial and temporal properties of the noise in a line-scan acquisition similar to the one used in Toi et al. measured from the human brain using a 3T MRI scanner. When measuring from the brain, we found temporal modulations with an amplitude spectrum that drops off as 1/f. These modulations were found in voxels spread across the reconstructed image along the phase-encoding direction. These artifacts were not present when scanning from a phantom. We built a simulation that replicates the qualitative and quantitative noise profiles observed in the data by breaking the assumption of stationarity of the underlying image between line acquisitions. These simulations suggest that the spatial spreading of temporal signals we observed in the human data is also due to nonstationarities between line acquisitions. In light of these results, we quantify how well one might be able to detect neural responses under different conditions and show that a low-amplitude neural response would be easier found as frequency response, rather than as a single peak. We end by discussing the issue of nonstationarity of neural responses and how this issue might present itself when trying to measure a DIANA response.

## Methods

### Participants

5 participants (3 male, 2 female) aged 20-35, naive to the goals of the study, participated in scans at the Stanford Center for Neurobiological Imaging. All protocols were approved beforehand by the Institutional Review Board for Human Subjects Research at Stanford University. All observers gave informed consent prior to the start of the experiment. Data for each subject were collected in 1-hour sessions.

### Scanner data acquisition

We collected data on a GE 3T UHP MR scanner with a 32-channel Nova head coil. Line-scan time series data were collected using a modified Cine sequence (Bohning et al., 1990; Lee et al., 2021; Waterton et al., 1985). An advantage of this type of sequence is data is acquired continuously so there is no disturbance in the steady state MRI signal, with cardiac gating applied retrospectively by sorting the data acquisitions according to a cardiac trigger signal. We used the same sequence parameters for both human and phantom data acquisition: a TR of 3.3 ms, TE of 1.5 ms, flip angle of 5 degrees, and bandwidth of 31.25 kHz. We collected voxels that were 3.75 x 3.75 mm in-plane and 5 mm through-plane. A single slice of 64 x 64 voxels is acquired each scan. Images were reconstructed from raw k-space data using an offline reconstruction for each TR, unlike the online GE reconstruction which temporally interpolates the line scan data to an equispaced number of timepoints (phases) throughout the cardiac cycle. We did not perform any post-processing on the data, including filtering, smoothing, or motion compensation.

We planned our slices with the goal of maximizing surface area of early visual cortex, so that we could characterize noise in sensory areas often used to validate MRI measurements and where one might expect to see a strong visual response to simple stimuli (Himmelberg et al., 2022). We collected T1-weighted images for each subject which we used to inform slice placement in the line scans. Coronal line-scan slices were placed by hand in posterior occipital cortex in an attempt to maximize the surface area of V1-V3. The T1-weighted image was also used for segmentation and surface reconstruction, which were performed using Freesurfer (Fischl, 2012). We aligned these reconstructions to a reference frame which we used to probabilistically predict early visual areas which we used in our SNR calculations (Benson and Winawer, 2018).

We collected three types of runs in which we varied synchronizing the start of each line acquisition to the cardiac cycle to determine whether this gating could reduce physiological noise. Each subject collected 8-10 runs of line scans in each of the 3 different conditions: one in which the start of each line acquisition was gated to the cardiac cycle (data acquisition begins at the peak of a photoplethysmographic waveform), one in which the line acquisitions were continuous (ungated by virtue of using an emulated cardiac trigger signal), and another ungated condition in which we swapped the frequency and phase encoding directions during image collection. Subject 5 did not collect data in the swapped phase-encoding condition. Because some of the scans were gated to the cardiac cycle, line duration varied for each subject based on the length of their cardiac cycle. Line duration is defined as the duration it takes to collect N repeats of one k-space line, where N is the desired number of reconstructed images in the time series. We set the line duration as equal to roughly 85% of the length of the cardiac cycle for each subject and kept it equal across all 3 conditions. In the cardiac gated trials, individual lines were resampled if the cardiac cycle to which it was gated was more than 20% faster or slower than the previous cycle.

In an exploratory run for one subject, we showed a stimulus that we knew would evoke a large neural response to test whether we might be able to measure said response. The stimulus was a full-contrast black-and-white checkerboard with a 1 cycle/degree spatial frequency that subtended -10 to +10 degrees of visual angle. The checkerboard appeared 50 ms after the start of each new line acquisition, contrast reversed at 300 ms, and disappeared at 550 ms. Stimuli were generated using MGL (Gardner et al., 2018) and Matlab and shown on a 60 Hz monitor. In all other acquisitions, subjects were not shown any stimuli and were instructed to remain awake with their eyes closed for the duration of the scans. All subjects were experienced with the scanner and had partaken in previous psychophysical experiments.

We also collected data from an fBIRN agar phantom (Friedman and Glover, 2006). We used the same acquisition parameters as in the human scans and placed the coronal acquisition slice in the middle of the phantom. The data was collected continuously (ungated) with a line duration of 825 ms (250 reconstructed images). We collected 3 scans of the phantom.

### Temporal structure: autocorrelation and frequency analysis

To quantify the amount of temporal structure in the data, we computed an autocorrelation permutation metric (APM). To compute the metric for each time series, we calculated the autocorrelation values of the time series: the correlation of the time-series with itself at all possible time lags. We then calculated a null autocorrelation distribution for comparison which specifies the distribution of correlation values expected for any time lag of a randomly organized (temporally unstructured) time-series by following the same procedure for 100 randomly permuted versions of the same time series. Permuting the time series lets us use the same samples but in random order, changing only the temporal structure. The autocorrelation metric for the time series is reported as the percentage of lags at which the non-permuted cross-correlation value falls outside of the 95% bounds of the permuted distribution for the same lag. This bound is two-tailed, as correlations can be positive and negative. This simple metric is used to characterize relative levels of temporal structure between voxels rather than as a significance test.

To determine the temporal frequencies in the data, we averaged the amplitude spectrum of all voxels that met an autocorrelation cutoff for each subject (> 0.70), and then calculated the spectral components using a Fast Fourier Transform. We then averaged the first 75 frequency components across subjects, a cutoff determined by the length of the shortest line duration of all the subjects. We fit curves to describe the amplitude distribution as a function of frequency of the form:

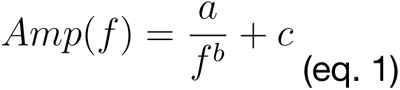

where a is a scaling factor, b is the exponential of the relationship between frequency and amplitude, and c is an offset. We used an adjusted r-squared to evaluate the fit of the curve that penalizes the fit for the number of parameters.

### Simulation

Our simulation was designed to reconstruct images from a ground-truth time-series of a modulating 2D image in a way that mimics how a line-scan acquisition reconstructs images. We first generate a ground truth image time-series of simple form: a square image with a square “brain” in the center that modulates with set frequency and amplitude. All other voxels are set to 0. This modulation is meant to mimic modulation in T2* contrast in the human brain from any mechanism. In the ground-truth sequence, the “brain” is a 16x16 square of voxels centered in a 65x65 image and each voxel has an average value of 100 throughout the simulation. All non-brain voxels have values of 0 throughout the entire simulation.

As this simulated brain modulates over time, we reconstruct images in a similar manner to how the line-scan acquisition builds out k-space, by building up lines of the 2-dimensional Fourier transform of subsequent images one line at a time (Fig 7). Our simulation reconstructs a sequence of images using similar parameters to those used in the actual data acquisition. In all simulations, we use a 1,000 ms line duration, 5 ms sample rate (TR), and sample 65 lines of k-space to reconstruct a 65 x 65 x 200 sequence (x, y, t). We use 65 lines rather than the 64 used in human data acquisition, as using an odd number of lines is convenient for examining the DC component as well as ensuring that all positive and negative frequency components are complex conjugate pairs. This conjugate pairing helps implement the half-Fourier approach described below. For each sample, we take the 2-dimensional Fourier transform of the image and add one line to the corresponding k-space image. We simulate each millisecond as 1 frame of the image sequence. Thus, the ground-truth sequence is 65,000 ms and is sampled every 5 frames. While generating unsampled timepoints is inefficient, it allows us to resample the same sequence with a different TR should we so desire. In the reconstruction, we implement a half-Fourier approach where instead of sampling all of k-space, we sample the first half and set the second half as the complex conjugate to ensure that our reconstructed images are real-valued. While, in practice, line-scan images contain complex-valued components that represent phase information, the results we present do not require simulating this phase component.

We simulated ongoing modulations in the brain voxels which, when reconstructed, produced effects similar to those observed in the data. We simulate this modulation either as a simple sinusoid (with a frequency, phase and amplitude) or as noise with a 1/f power spectrum (pink noise). In an ideal case the underlying modulation during each line acquisition is exactly the same. Breaking this assumption in our simulation replicates spatial properties of the noise in our data. That is, nonstationarity in the underlying image between line acquisitions due to sampling at different points in the underlying modulation spreads that modulation across the image.

For convenience, to induce nonstationarity between line acquisitions when modulating the brain voxels as a sinusoid, we randomized the phase of the sinusoid at the beginning of each line acquisition. This offsets the phase of the modulation at each line while modulating at an integer frequency, which would otherwise not be offset as it would perfectly divide the 1-second line duration. Modulating at an integer frequency allows us to look directly at the phase and coherence maps of the modulation frequency in the reconstructed data. We cannot do the same for non-integer frequencies (e.g. we cannot look at the 2.7th frequency component). It is worth noting that both approaches to induce nonstationarities between lines – randomizing the phase of an integer frequency at each line or modulating at a non-integer frequency – produce similar results. To induce nonstationarity between lines in the pink noise modulation case, we generate independent time series for each line acquisition with the same amplitude spectrum.

We also examined the effect of modulations which were not globally coherent across the image. In these cases, instead of modulating all voxels in the simulated brain together, we modulated spatial groupings of voxels together with offsets between the groups. In the sinusoidal modulation case we added unique phase offsets to each group throughout the entire simulation. To define the groups, we split the brain into equally sized rectangles along the phase-encoding direction or into squares (groups and example offsets pictured in figure 9).

### Noise and Signal-To-Noise Estimates

We examined the effect of averaging across voxels on the magnitude of the noise. It is almost always the case that experiments will average a response over contiguous voxels, for example over a sensory area. Thus, we calculated each voxel’s (all voxels from areas V1-V3) Euclidean distance from the center of mass of that cluster of voxels and added them one-by-one by distance to an averaged time series, calculating the standard deviation of that time series at each step. We fit this relationship – between the number of voxels averaged and the standard deviation of the time series – with an exponential function of the form:

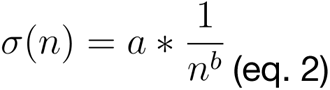

Where sigma is the standard deviation of the time series as a function of the number of voxels n, and “a” was fixed as the average standard deviation of the time series of all of the individual voxels in V1-V3. “b” was either fit to the data or fixed as a specific exponent. In the case of completely independent voxels, we expect a square-root-of-n (b = 0.5) reduction in the noise magnitude. We compared the observed noise reduction in each scan to this expectation by taking the ratio of equation 2 with the fit b value to equation 2 with b = 0.5, evaluated at n equals the number of V1-V3 voxels in each subject.

We performed a similar procedure to calculate the effect of averaging over multiple scans on reducing noise. For each scan, we averaged all V1-V3 voxel time series together. Then, for each subject, we averaged over an increasing number of scans similar to how we averaged over an increasing number of voxels and measured the effect on the standard deviation of the time series. Because some scans are noisier than others, for each number of scan averages n, we sampled every possible combination of n scans and took the average standard deviation of those averaged time series. Again, we fit an exponential function (eq. 2) to quantify the relationship between number of scans averaged and magnitude of the noise and compared it to a square-root-of-n noise reduction (b = 0.5), which is expected from independent scans. Here, “a” is fixed as the average standard deviation of all individual scan time series for each subject and the ratio is evaluated at n equals the number of scans collected.

We used these measures of how averaging over voxels and scans reduces the magnitude of the noise to estimate our ability to detect a neural response of specific magnitude in the presence of the noise we measured in our scans. We used empirical measurements to estimate the expected noise level under different conditions. In the impulse response condition, we then calculate the signal-to-noise-ratio (SNR) as ratio of the magnitude of the expected response to the magnitude of the noise.

To make SNR estimates, we needed estimates of the response and noise magnitudes. Here we assume an expected response to a stimulus of 0.2%, which is similar to the strongest responses described in Toi et al. To estimate the noise level as a function of how many voxels were included, we repeated the procedure described earlier in which we fit an exponential function (eq. 2) to describe the noise magnitude as a function of voxels included. Here we fit the function to all data across all subjects, instead of to individual subjects. This provides an estimate of the noise level with different numbers of voxels for a generic subject. Based on our previous analysis, we approximated the reduction in noise as a function of averaging scans as a square-root-n reduction. Together, these estimates give us a noise magnitude estimate based on the number of voxels and scans, which we can use to calculate SNR given the stimulus response estimate. We defined SNR in the impulse response case as the peak of the expected response (0.2%) divided by the standard deviation of the noise.

Because the expected response is relatively small, and because the noise in our data drops off at high frequencies, we hypothesized that it might be easier to detect neural responses that modulate at a single frequency over time. Toi et al. measured the neural response following a brief stimulus presentation, also known as an impulse response. Another approach to measuring a visual response is to modulate a visual stimulus over time and measure the response at the stimulus frequency of its harmonics (Regan.1966, Engel.Shadlen.1994, Boynton.Heeger.2012, Norcia.Rossion.2015). In this frequency response approach, the stimulus is modulated at a frequency outside of the dominant noise frequencies, making the neural response easier to detect. To compare this frequency approach to the impulse approach outlined above, we drew a similar SNR line as a function of scan averaging. This was done via simulation; first, we simulated 1/f noise of the same magnitude as the average scan across all subjects. We then added a sinusoidal response of the expected magnitude (0.2% signal change from peak to trough) at 5hz to the time series. To estimate the noise at the stimulus frequency, we took the average of the four adjacent frequency component magnitudes (-2 to +2 cycles/second) from the simulated time series. SNR was then defined as the ratio of the magnitude of the response frequency (minus the noise estimate) to this noise estimate. We averaged this value across 1,000 simulations. We repeated this procedure across different numbers of scan averages, meaning that we reduced the 1/f noise amplitude as a square root of the theoretical number of scans included.

## Results

### Initial measurements in visual cortex: false positive responses

We collected line-scan data from a participant who viewed a large, high-contrast checkerboard stimulus. The stimulus was presented 50 ms after the start of each line acquisition. The stimulus contrast-reversed 250 ms after first appearing and disappeared 250 ms later. This stimulus evokes a large neural response in visual cortex (Engel et al., 1994; Gardner et al., 2008). Averaging over voxels in V1-V3, across 10 scans with identical protocol, we measured a signal comprising two peaks, roughly aligned with stimulus onset and contrast reversal (Fig 1a), that could easily be interpreted as a neural signal.

**Figure 1:**
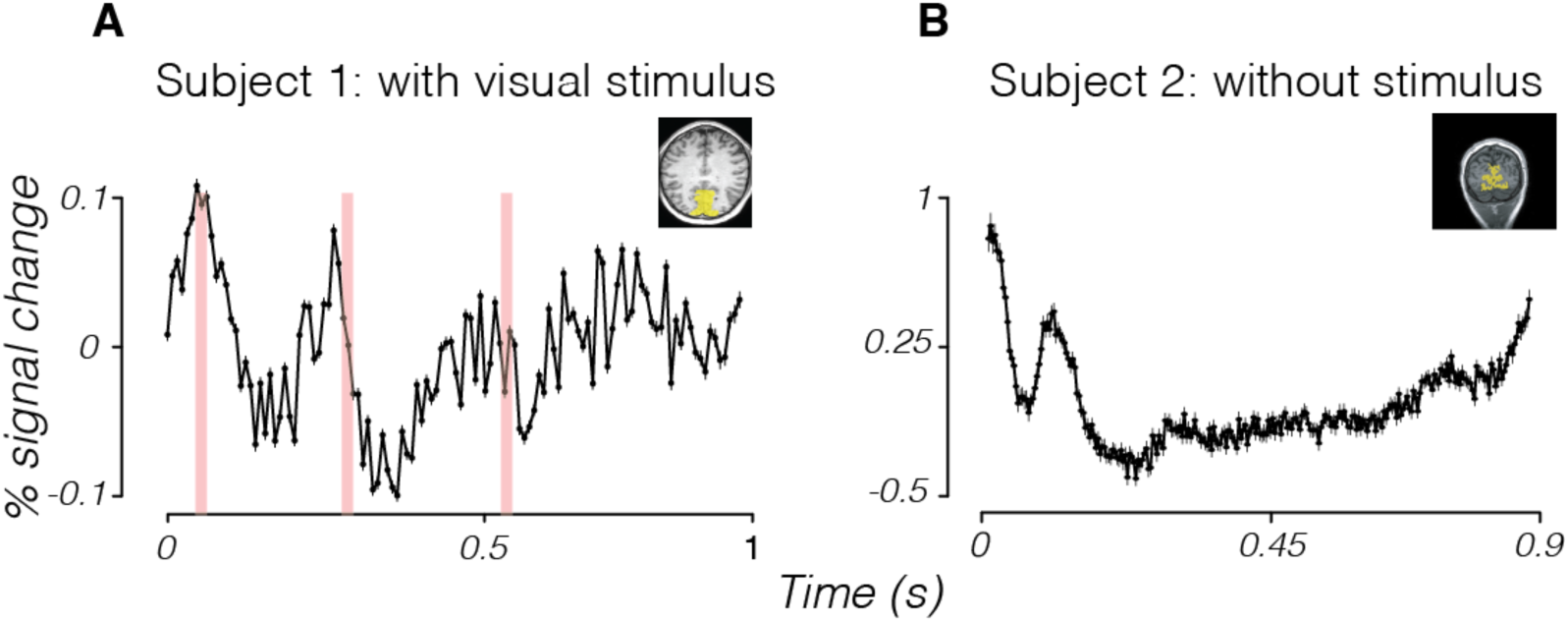
MR line-scanning time series in human visual cortex comparing a stimulus and no-stimulus condition. Time series for voxels averaged across V1-V3 (yellow shading in insets showing T1-weighted slice image). (A) Time series from a subject viewing a flickering stimulus comprising a black-and-white checkerboard with an onset at 50 ms, a contrast-reversal at 300 ms, and offset at 550 ms (times marked in red). (B) Time series from a subject with eyes closed and no stimulus. The modulations here are of greater magnitude than when a stimulus was shown.

We ran the same sequence on a second subject without a visual stimulus to ensure that the peaks we observed were due to a visual response. This control condition also had a large peak in the data that could easily be construed as a neural response (Fig 1b, also averaged across V1-V3 and 10 scans). For technical reasons these data were collected with a shorter TR, resulting in finer sampling. This result shows that noise in the line-scan acquisition contains fluctuations whose temporal structure can be similar to the expected signal and are different from random Gaussian noise.

### Temporal characteristics: Low Frequency Modulations

### Phantom

To further characterize the noise and investigate the potential source of the noise, we acquired line-scan data from a slice of an agar phantom (Fig 2a). We collected and averaged 3 scans using parameters similar to Toi et al. (TR of 3.3 ms, TE of 1.5 ms, and line duration of 825 ms, 250 reconstructed images). To quantify the overall noise level we computed the mean and standard deviation of the signal averaged across all voxels in the phantom (Fig 2b). The time series offers a baseline measurement of the noise level. Averaged across all voxels (n = 1383), the standard deviation of all time points was 0.014% of the mean.

**Figure 2:**
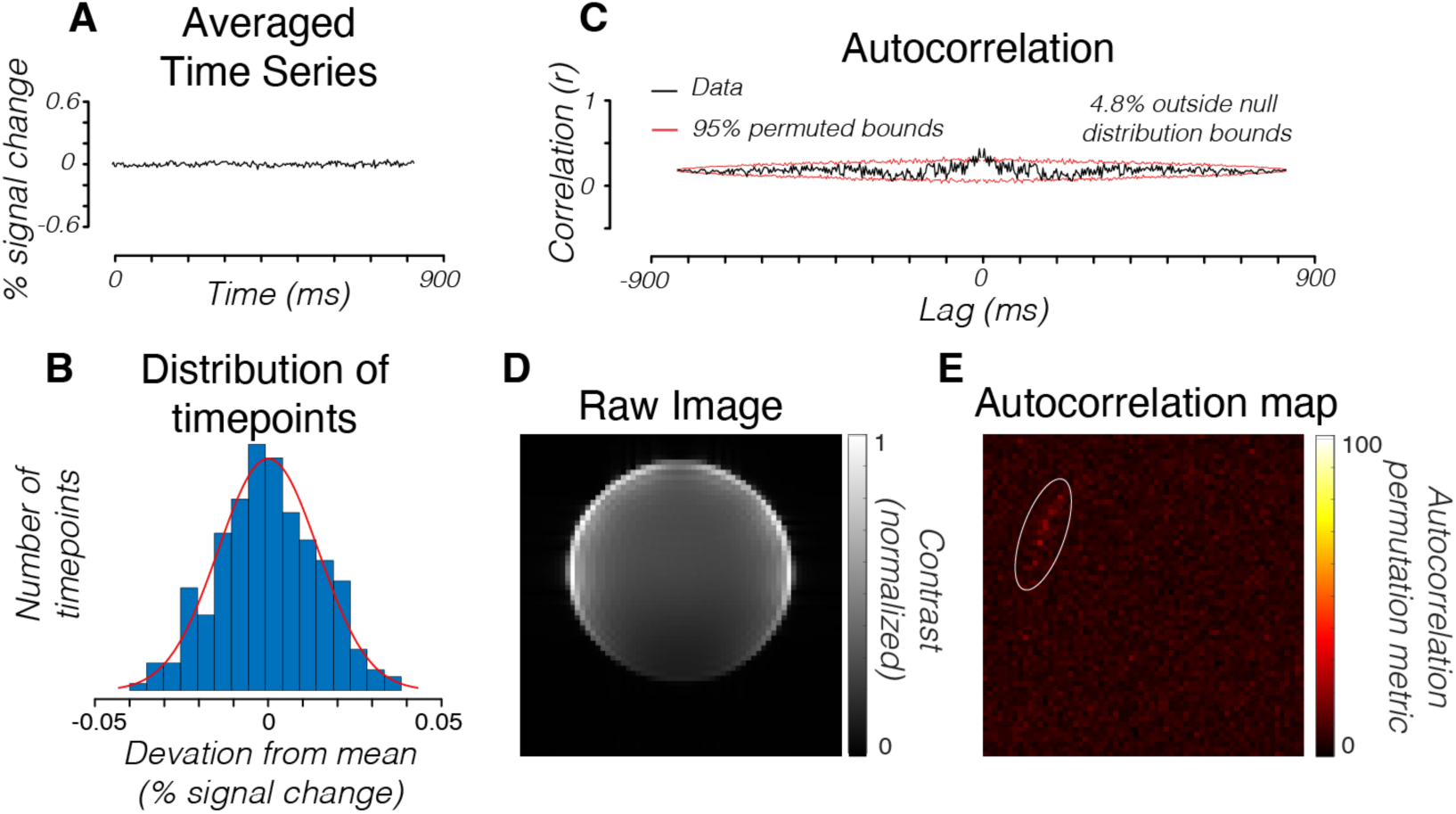
MR time series from a phantom using the line scanning protocol. (A) Average time series measured in all voxels (n = 1383) in a middle-slice of an agar phantom across 3 repetitions. (B) Histogram of percent signal change from the mean of the data. Each observation is a single time point in the time series. Red curve is a Gaussian distribution fit. (C) The autocorrelation permutation metric (APM) of the time series averaged across all voxels in the phantom. Red curves bound the confidence interval that contains 95% of the autocorrelation values computed after temporally shuffling (randomizing) the time-series (100 repeats). In the phantom time series, 4.8% of the time points were outside of the 95% permuted bounds. (D) Mean intensity of each voxel averaged across all time points in the scan. (E) A map of the APM metric for each voxel. In the phantom the noise is approximately Gaussian, independent, and identically distributed. The non-zero autocorrelation is very small, and appears to be due to pixels at the edge of the phantom (white circle). This may be due to small movements during the acquisition due to scanner vibrations.

To quantify the temporal structure of the noise, we computed the autocorrelation of the time series and compared the amplitude of this autocorrelation with the expected autocorrelation amplitude when the time series was randomly shuffled, removing any temporal structure (red lines in Fig 2c, full details in Methods: Temporal Structure). The fraction of time points at which the autocorrelation amplitude exceeds the shuffled autocorrelation amplitude is reported as the autocorrelation permutation metric (APM). This metric quantifies the temporal structure in the time series. Were the noise perfectly independent and identically distributed, the autocorrelation amplitude would be constant and the APM zero. We observed only a small deviation from this ideal in the phantom averaged across all voxels (Fig 2c, 4.8% APM). Individual voxels in the phantom also had low APMs (Fig 2e, mean = 6.03%, std = 2.01%, n = 1383 individual voxels).

### Visual cortex

To characterize the noise in humans, we acquired line-scan data from 5 subjects using the same parameters as the phantom. Subjects were not shown any visual stimuli and were instructed to remain awake with their eyes open for the duration of the scans. Each subject collected 8-10 scans of data (details in Methods: scanner data acquisition). For each subject, we averaged together all scans and selected a region of interest in visual cortex covering V1-V3 over which we averaged voxel time series. These human-subject time series had significantly higher standard deviations than the agar phantom, even when averaged over more scans (std = 0.17%, 0.26%, 0.24%, 0.19%, 0.22%, compared to 0.014% signal change in the phantom, p < 0.001 for all subjects, one-sided t-test).

Unlike measurements from the phantom, the human brain time-series had substantial temporal structure when averaged over areas V1-V3 (Fig 3a), as evidenced by high autocorrelation permutation metrics (Fig 3b). This temporal structure was apparent when averaging the time series across voxels in early visual cortex (V1-V3) in each subject (Fig 3b: Mean = 65.8%, std = 15.1% across 5 subjects). The temporal structure is also evident in individual voxels in the whole brain (Fig 4b,e) (mean APM = 62.42%, std = 17.67%, n = 1384 total brain voxels across 5 subjects).

**Figure 3:**
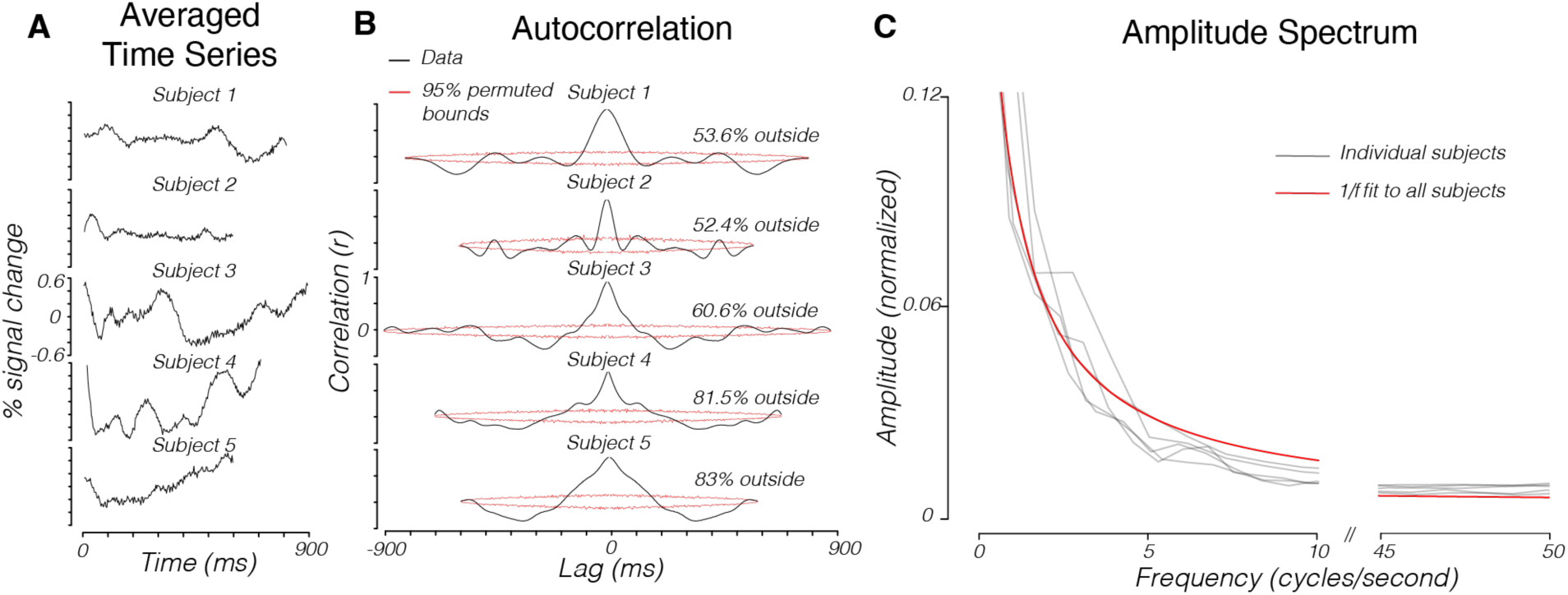
The noise distribution of MR line-scan time series in human visual cortex. (A) The MR line-scan responses for 5 human subjects. Each curve is averaged over V1-V3 (65-116 voxels, depending on subject) across 8-10 repetitions. Slices were acquired from posterior occipital cortex. (B) Autocorrelation permutation metrics for each time series. (C) Averaged, normalized amplitude spectrums of voxels with APM values greater than 0.7. Gray lines are individual subject data. Red line is a 1/f fit to all subjects’ data (equation 1, methods).

**Figure 4:**
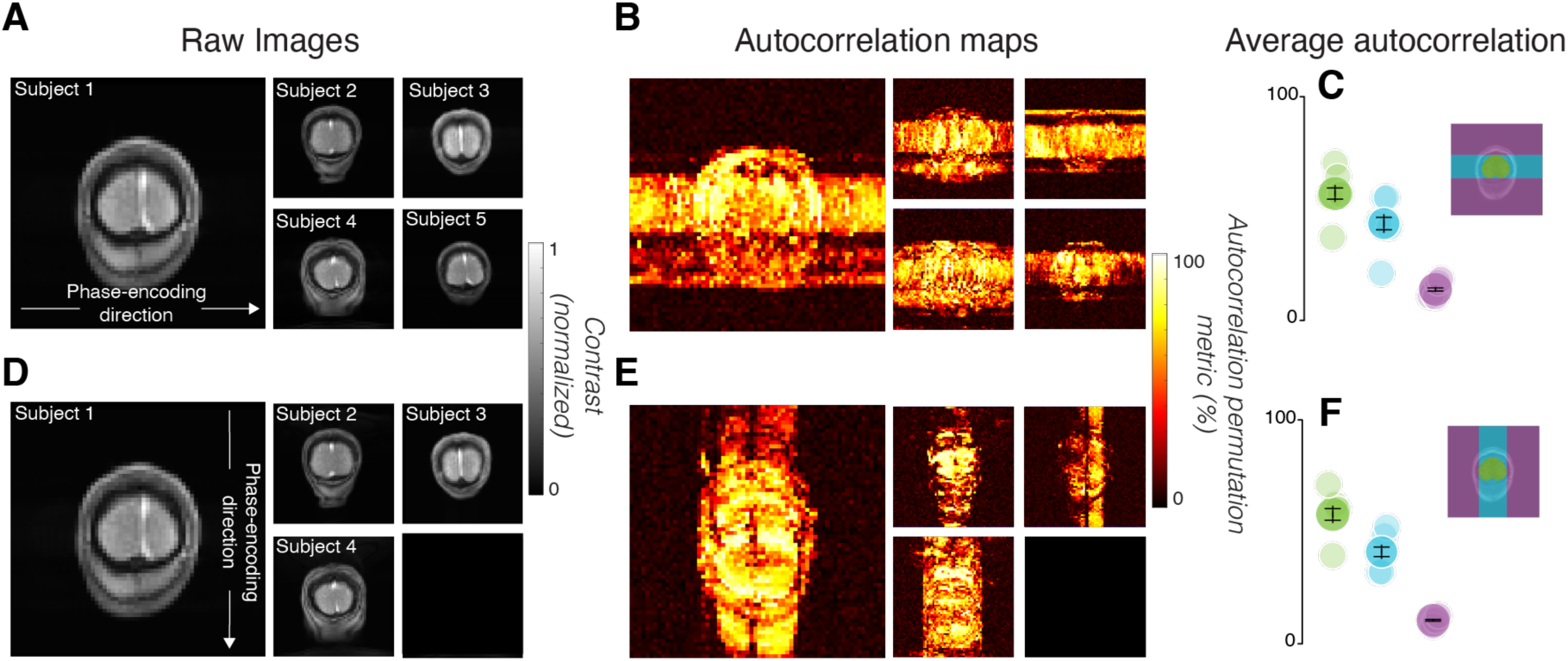
Autocorrelation maps depend on the phase-encoding direction of image acquisition. (A) Raw images acquired with the line-scan sequence for each subject, averaged over 8-10 scans and averaged across time. The phase-encoding direction was along the x-axis of the image. (B) Maps of the autocorrelation permutation metric. (C) The average autocorrelation permutation metric from In-Brain (green), Outside-Along (cyan) and Outside-Against voxels (purple), segmented for each subject (one example shown). Lighter points are individual subject data; darker points with error bars are averages of all subjects. Error bars are 1 standard error of the mean. (D-F) Same as A-C, but with the phase-encoding direction along the y-axis. Data for subject 5 are missing in this condition due to technical error.

We explored the time series of all voxels with robust temporal structure (high APM values) and found that their time series have a 1/f amplitude drop-off (Fig 3c). For each subject we normalized and averaged the amplitude spectrums of the voxels with high autocorrelation permutation metrics (> 0.70). The amplitude spectrum was well-described as 1/f (methods, equation 1) (Fig 3c, adjusted r-squared = 0.94).

### Modulations are spread along the phase-encoding direction

Voxels with high autocorrelation metrics were found in the brain, in the skull, and in the empty volume outside the head. Many voxels that contained a segment of the brain within its phase-encoding line had high autocorrelation permutation metrics, even those outside of the head (Fig 4b). Given this finding, we labeled voxels outside the brain along phase-encoding lines that include the brain as ‘Outside-Along’; voxels on PE lines that do not include the brain are referred to as ‘Outside-Against’; voxels inside the brain are referred to as ‘In-Brain’ (cyan, purple, and green regions in Fig 4c). We identified and labeled these voxels by inspecting the T1-weighted image (Fig 4a). The mean APM for In-Brain voxels was higher than the mean APM for Outside-Against voxels (Fig 4b,c, p < 0.001, one-tailed t-test comparing green to purple points, mean APM = 62.15% vs. 15.28%). The mean APM for Outside-Along voxels was also higher than for Outside-Against voxels (Fig 4b,c, p < 0.001, one-tailed t-test comparing cyan to purple points, mean APM = 53.95% vs 15.28%).

To test whether the spatial spread of temporal structure in the time series was dependent on the positioning of the phase-encoding direction, we swapped the phase- and frequency-encoding (readout) directions during image acquisition. The new Outside-Along voxels once again had a higher APM than the new Outside-Against voxels (Fig 4e,f, p < 0.001, one-tailed t-test comparing cyan to purple points, mean APM = 39.76% vs. 9.37%).

Given the amplitude spectrum of the time series in high APM voxels, it is also possible to summarize the spatial spread using the amplitude of the lowest temporal frequency component. The image based on the coherence (amplitude at f divided by the sum of amplitudes across f) (Fig 5a) matches the image based on the APM (Fig 4a) (r = 0.93, 0.80, 0.91, 0.92, 0.86, pointwise Pearson’s correlation between coherence and APM maps). Thus, we can explore the spatial spread of the signal by examining both the amplitude and phase of this first frequency component (Fig 5b).

**Figure 5:**
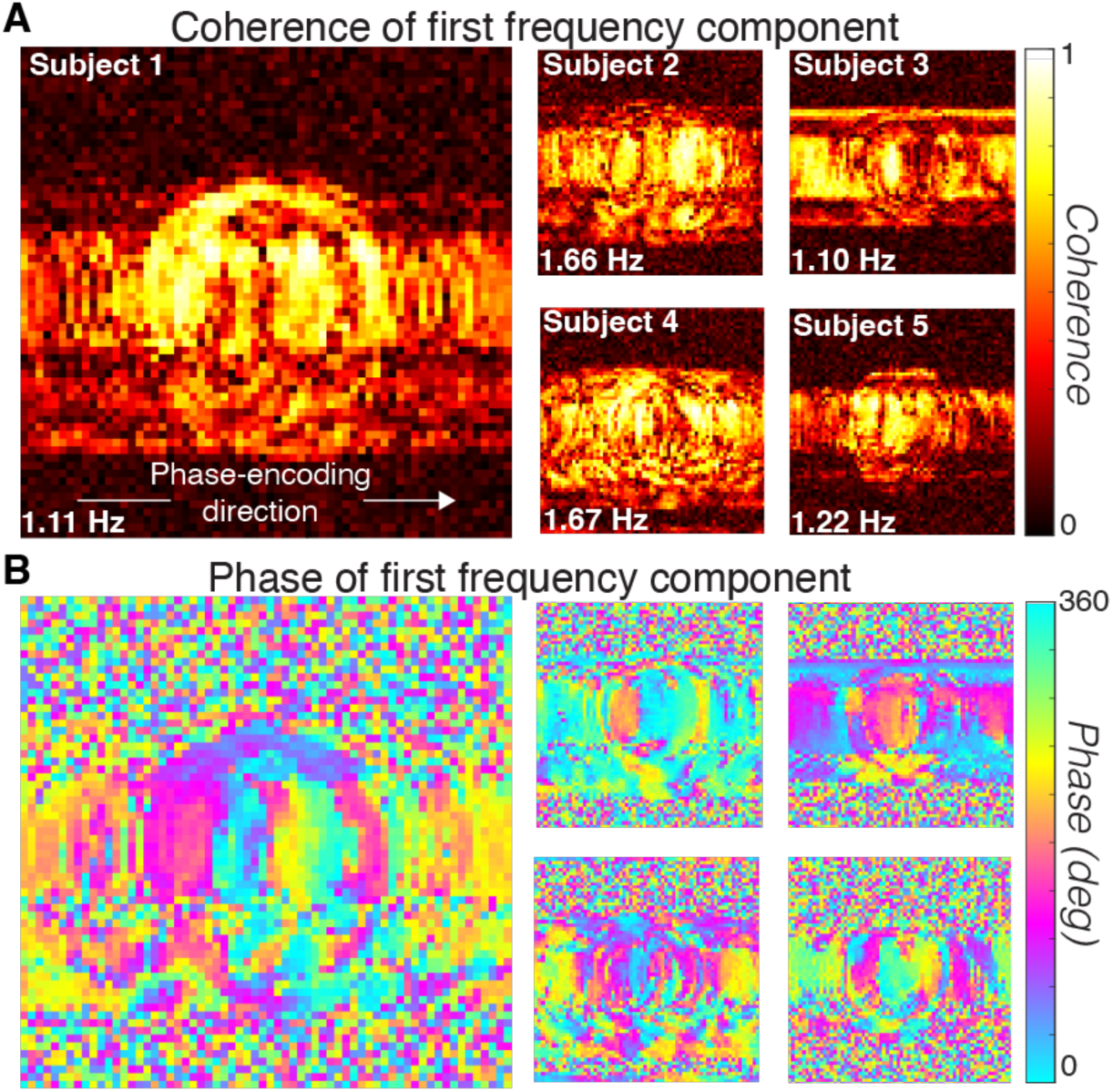
Spatial structure of the first frequency component coherence and phase. The noise is greatest in the low temporal frequency terms (Fig 3c). Maps show the coherence (A) and phase (B) of the lowest temporal frequency (see image text). Coherence is the amplitude of the lowest temporal frequency divided by the sum of the amplitudes of all frequencies. The coherence is very low in the Outside-Against region and the phase is random across space. Within the In-Brain and Outside-Along voxels the phase pattern for each subject is robust and approximately constant perpendicular to the phase-encoding direction.

The phase of the first frequency component reveals timing differences in the modulations across the image. The phase of the first component varies smoothly along the phase-encoding direction (Fig 5b) across the In-Brain and Outside-Along regions.

The phase is nearly constant along the direction perpendicular to the phase-encoding direction. The phase has no obvious spatial structure in the Outside-Against regions. This variation suggests that temporal modulations within the brain introduce systematic timing differences in nearby regions that spread in the phase-encoding direction. We do not see this pattern in the phantom because there are no corresponding temporal modulations.

### Cardiac gating did not eliminate the spatial spread

The heartbeat is one potential source of systematic temporal modulations during the acquisition, the timing of which may differ between line acquisitions. To reduce the variations on the line scans with respect to the heartbeat, we gated the start of each line acquisition to the heartbeat while collecting data.

Gating the start of each line acquisition to the heartbeat did not significantly change the autocorrelation permutation metrics in the time series inside or outside of the brain. For each subject, the average APM of voxels in different regions of interest were little changed when comparing the gated and ungated conditions (Fig 6). In the cardiac-gated acquisition, In-Brain voxels (Fig 6, left, green vs. blue, p < 0.01, paired t-test, n = 5) and Outside-Along (Fig 6, left, cyan vs. purple, p < 0.01) had higher APM levels than Outside-Against voxels (one-tailed t-tests across 5 subjects). However, there was no significant difference between the APM levels in the gated and ungated conditions in In-Brain voxels (Fig 6, green points, 0.56 +/- 0.13 std vs. 0.62 +/- 0.03 std, p = 0.43, paired t-test, n = 5), Outside-Along voxels (cyan points, 0.44 +/- 0.15 std vs. 0.54 +/- 0.09 std, p = 0.16), or Outside-Against voxels (purple points, 0.14 +/- 0.03 std vs. 0.15 +/- 0.05 std, p = 0.38).

**Figure 6:**
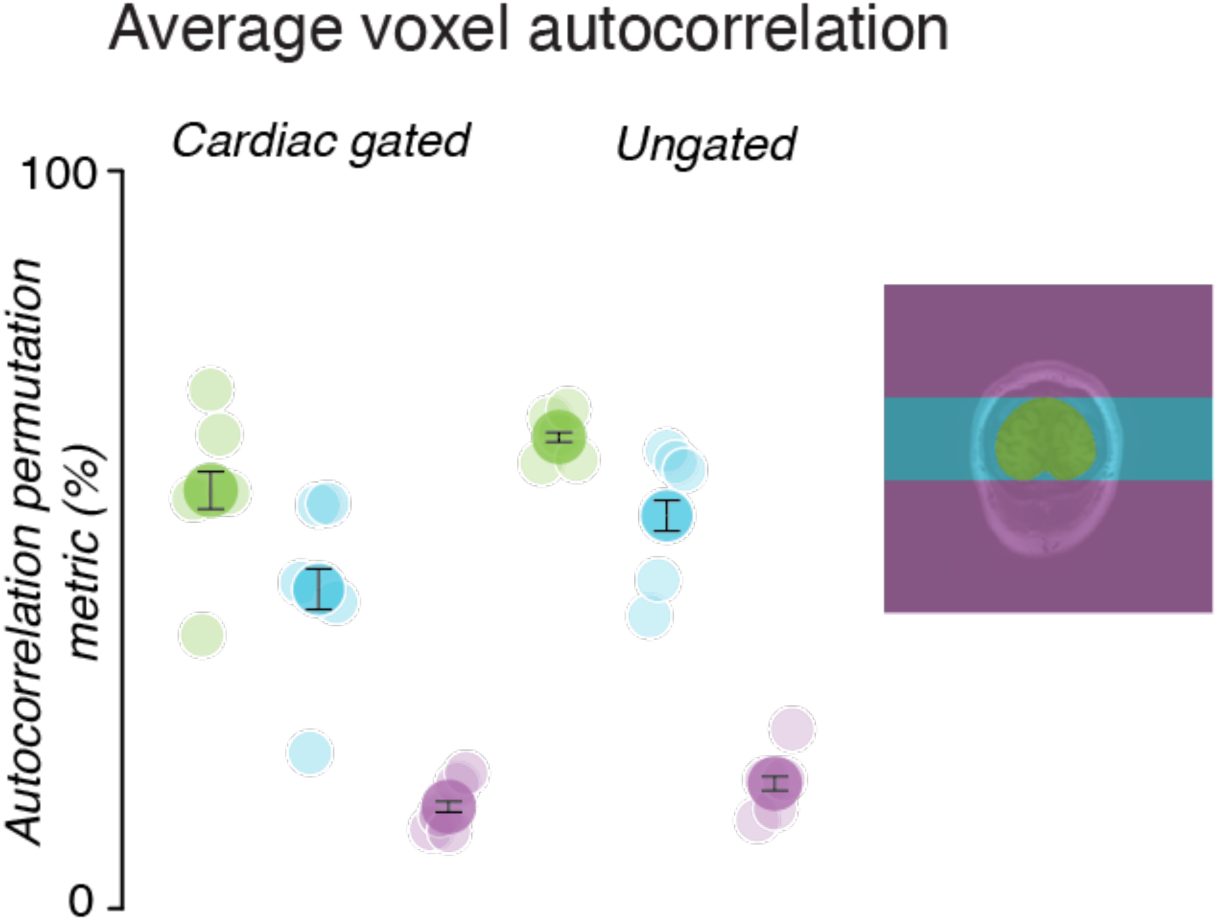
Autocorrelation permutation metrics are similar when using cardiac-gated and free running conditions. The average APM across all subjects, along with the SEM, are shown by the darker points. Lighter points show the average APM for each subject. The colors correspond to different regions (image at right).

## Model & Simulations

The spatial and temporal structure of the noise could arise from nonstationarities of the imaging substrate during image acquisition. In line-scan acquisitions, the impact of nonstationarities differs between the two dimensions of k-space. In the fast direction – the frequency-encoding direction – the k-space samples are acquired over a few milliseconds. In the slow direction – the phase-encoding direction – the k-space samples are acquired up to a minute apart. The brain substrate is far less likely to remain in the same state across k-samples in the slow direction.

We simulated the impact of this interaction between substrate modulations and the k-space line scan acquisition. The simulation generates a time-varying ground-truth image time series and reconstructs the time series expected from a line-scan acquisition (Fig 7). The modulating “brain” is modeled as a square group of pixels with a mean value of 100 that modulates over time as a simple sinusoid with a modulation amplitude of 1% (Fig 7a,b). In the simulation, the brain modulates at 3 Hz (the “modulation frequency”). The line scan protocol samples 65 lines. Each is sampled repeatedly every 5 ms over a 1 second duration. The total time for acquiring all lines is 65 seconds. This protocol generates an image sequence comprising 200 images sampled at 5 ms intervals with 65 by 65 spatial samples.

Acquiring each line of k-space at the exact same phase of the underlying modulation – the stationary case – recreates the image exactly. Acquiring different lines at different times with respect to the underlying modulation – the nonstationary case – means that k-space lines are intermingled that modulate with different phases (Fig 7b). The simulation purposely induces nonstationarity between line acquisitions by randomizing the phase of the sinusoidal modulation at the beginning of each line acquisition. The way we reconstruct images – by taking single lines of the Fourier transform of the ground-truth image – mimics the fast and slow acquisition dimensions in the line-scan acquisition. Each line is acquired during a time period in which the substrate is roughly constant; but, adjacent lines are collected at times when the simulated substrate level might be quite different. This procedure simulates the main properties of the MR line-scanning protocol.

### Model of line-scan acquisition recreates spatial spreading artifacts

The nonstationarity of the substrate between line acquisitions produces reconstructed images with spatial artifacts similar to the measurements (Fig 8). In the simulation, brain voxels modulated over time and the voxels outside the brain were fixed to zero. Despite this, the reconstructed time series using the line scan protocol had higher autocorrelation permutation levels in the Outside-Along voxels than the Outside-Against voxels (Fig 8b,c, voxels in cyan vs. purple areas, APM = 0.77 vs. 0.076, p < 0.001, one-tailed t-test). In the brain and Outside-Along voxels, the reconstructed response is at the modulation frequency (Fig 8d,e). The phase of the modulation frequency varies across the phase-encoding directions (Figure 8f), similar to the phase variation in the human data (Fig 5f).

**Figure 7:**
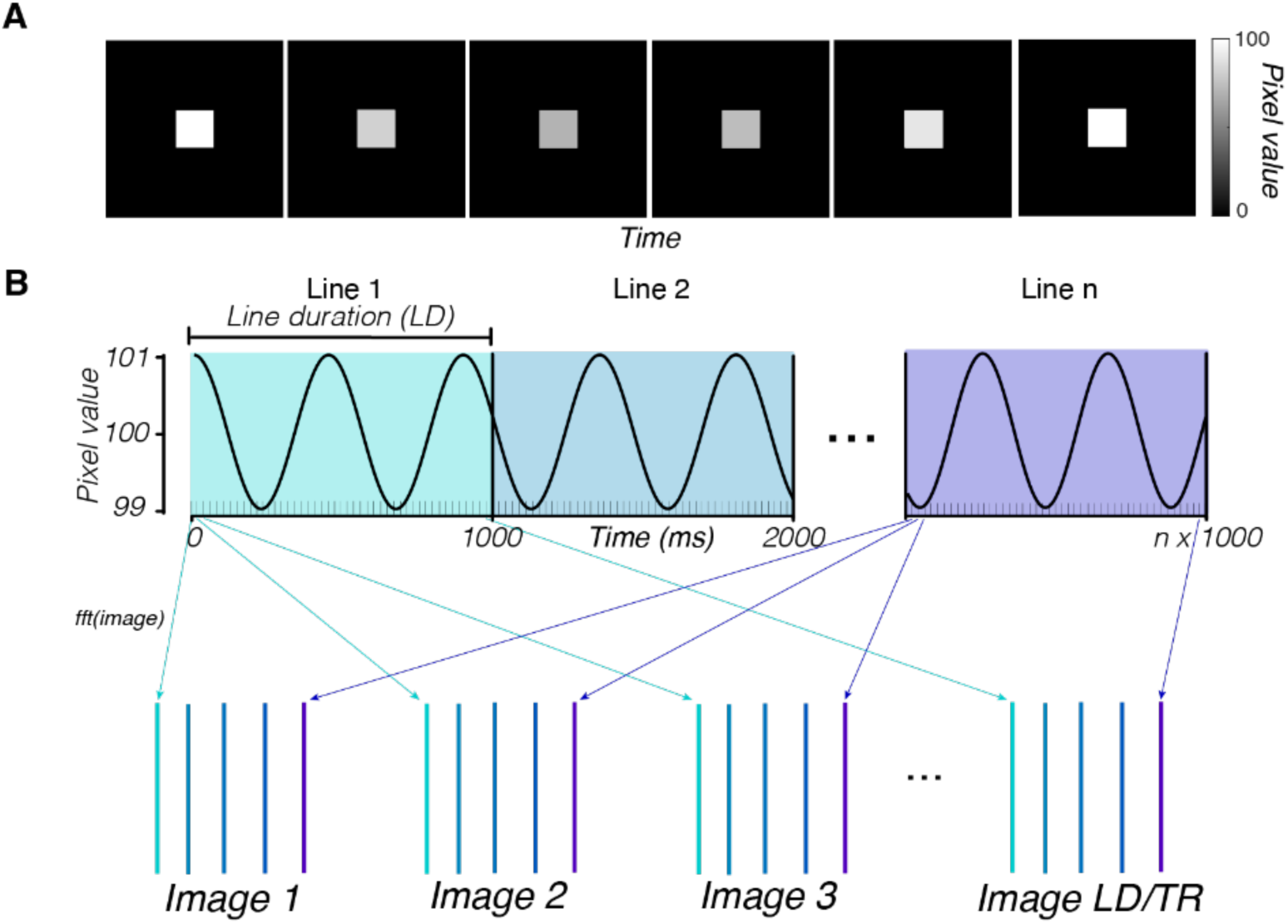
Simulation of a line-scan acquisition from a time-varying substrate. (A) The input substrate is modeled as time varying; its intensity modulates as a 3 Hz sinusoid around a mean (100, 1% modulation). Voxels outside the region of the square substrate are constant at zero. (B) Line acquisitions start at random times with respect to the sinusoidal modulation, creating nonstationarities between line acquisitions. Each line is sampled 200 times – once every 5 ms – over the course of a second. The figure illustrates how data from the first and last line acquisitions are used to reconstruct each of the images in the time series. With this procedure, each image includes data acquired over a full minute. After the full k-space images are acquired, we use the inverse Fourier transform to convert them to real images.

**Figure 8:**
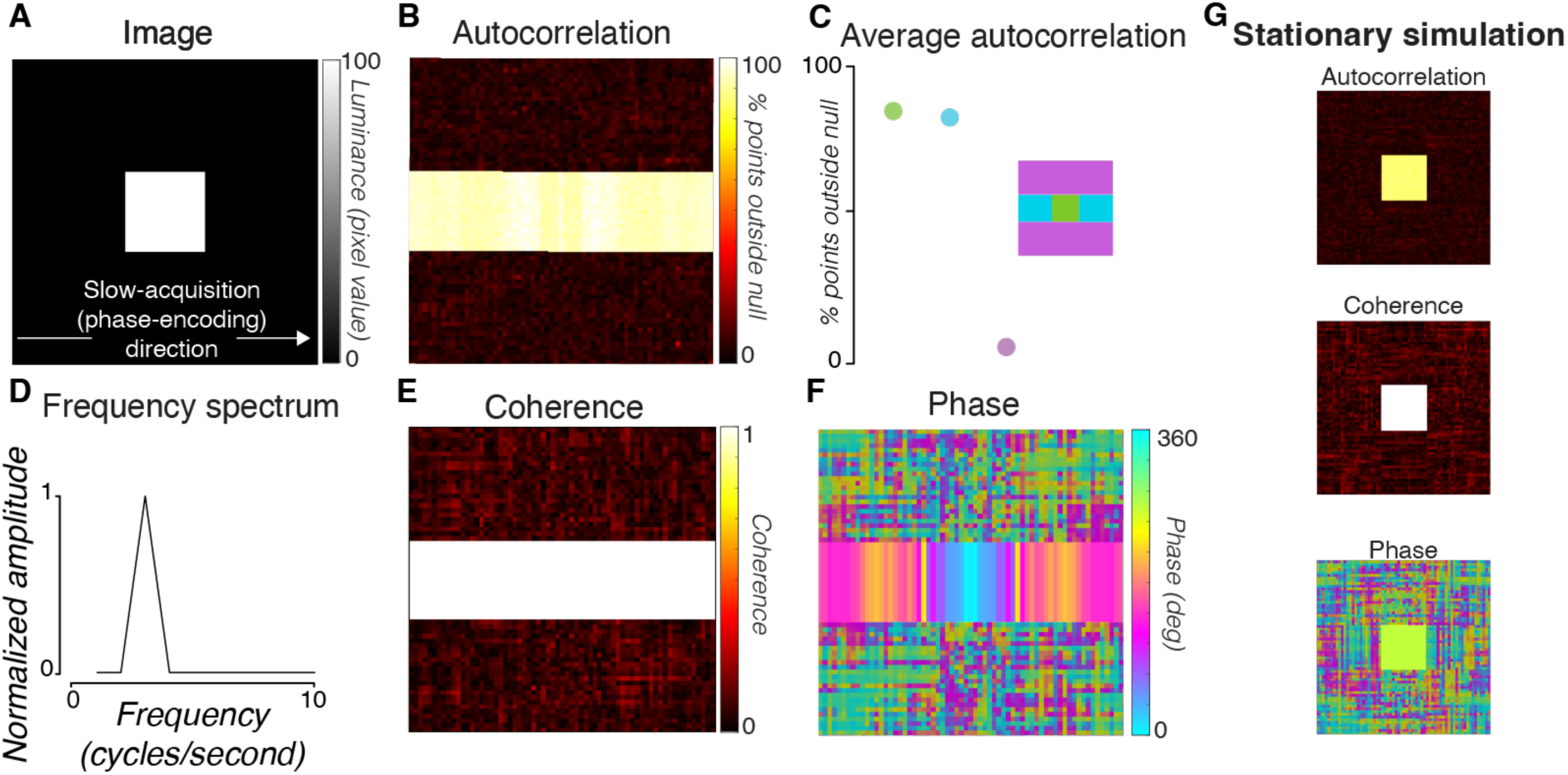
Simulation replicates spatial and temporal patterns in the data. The substrate was modulated at 3 Hz and each line scan was acquired at a random time during the modulation. (A) Reconstructed image, averaged across all time points. (B) A map of the autocorrelation permutation metric. (C) Comparison of the autocorrelation permutation metrics in different regions of the image, shown in the inset. (D) Normalized frequency spectrum averaged over all In-Brain and Outside-Along voxels (cyan + green areas). (E) Coherence map of the substrate modulation frequency. (F) Phase map of the same component. In the simulation phase maps we darkened the Outside-Against voxels, as their time series should be zero (any modulation is rounding error) and thus have indeterminate phases. (G) Same as B, E, F, but for a simulation in which each line scan was acquired at the same phase of the modulation.

**Figure 9:**
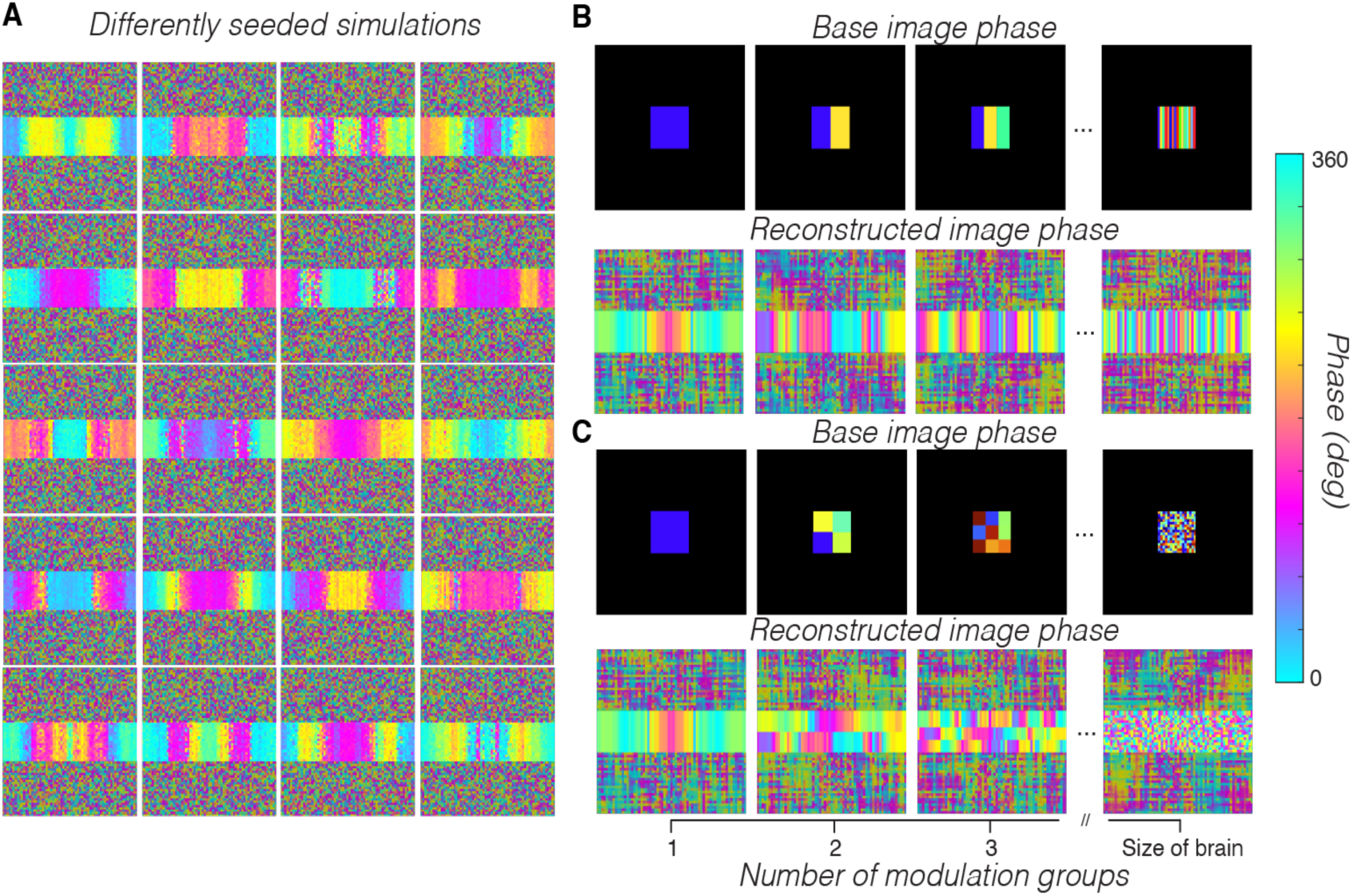
The phase map depends on the spatiotemporal distribution of the substrate modulation. (A) We simulated 20 time series, each from a 1/f noise distribution, and reconstructed the image time series using a line scan protocol. The maps show the phase of the first frequency component. The details of the map vary between different simulations, but all the maps show the same basic pattern of spreading from the brain along the phase-encoding direction. (B,C) We then returned to using a 3 Hz substrate modulation, but varied the phase of the modulation across spatial locations. The top rows show the phase of the simulated sinusoidal modulation. The bottom rows show the phase map of the first frequency in the reconstructed image series. The time series of the black voxels around the simulated substrate have zero mean and zero modulation.

If each line acquisition begins at the same phase of the substrate modulation, the spatial spread is eliminated. The large autocorrelation inside the simulated head remains, and the phase is constant across the substrate (Fig 8g). This observation motivated our experiment with cardiac gating, in which the gating did not reduce the spreading in practice. Note that the autocorrelation, coherence, and phase maps are symmetric across the phase-encoding direction in this simulation. This symmetry in the reconstructed images is due to the symmetry and centrality of the modulating substrate region as well as the globality of the modulations. The symmetry of the maps is broken by changing these assumptions (example illustrated in Fig 9b,c).

### Brain modulations influence the phase gradient

How the underlying modulation spreads across the image along the phase-encoding direction depends on the offset of the modulation between subsequent line acquisitions. To illustrate this dependence, we ran 20 differently seeded iterations of the simulation and examined the phase gradient that emerged (Fig 9a). Here, to simulate modulations closer to what we observed in the human brain, we modulated the ground-truth voxels globally with a 1/f drop-off in amplitude (pink noise) rather than as a sinusoid. This pink noise was regenerated with a random seed for each line acquisition, resulting in a different sequence of noise with the same amplitude spectrum for each line. In addition, we added a small amount of independent Gaussian noise to each ground-truth voxel time series. The phase maps of the first frequency component in each simulation show how the spread of the signal is dependent on the random offset between lines, with different randomizations producing very different patterns of spread.

To determine how differences in the spatial structure of the simulated modulation change the phase of the reconstructed modulations we ran simulations using a single sinusoidal frequency but randomizing the phase between different subregions. Instead of modulating the brain in the ground-truth time series with global coherence (all modulating voxels have the same time series), we systematically break that assumption by breaking up voxels in the brain into smaller and smaller modulation groups that are offset from each other, either as rectangles (Fig. 9b) or squares (Fig. 9c, “base image phase”).

Breaking the assumption of global coherence in the simulation produces a progressively less-ordered phase gradient of the major frequency component along the phase-encoding dimension of the reconstructed image (Fig. 9b,c, “reconstructed image phase”). These results show that varying how globally coherent the underlying modulation is in the simulation has systematic effects on the phase gradient of the underlying modulation. Conversely, this result suggests that the observed phase gradients in the human data (Fig. 5b) may be diagnostic of the underlying modulation that is causing spreading along the phase-encoding dimension. They suggest a strong, globally coherent underlying modulation in the brain and parts of the skull.

### Expected SNR across different stimulus types and scan- and spatial-averaging choices

We used our empirical noise measurements to estimate how SNR for the line-scanning technique would be expected to change for different amounts of trial (number of scans) and spatial (number of voxels) averaging, as well as under different modes of stimulus presentations. To do so, we calculated from our human measurements how the magnitude of the noise decreases as a function of averaging over an increasing number of scans. We calculated a similar relationship between the magnitude of the noise and the number of voxels averaged. Using these estimates of the magnitude of the noise, we calculate how many scans one must average to achieve specific SNR given the amount of cortex they are averaging over and the strength of the expected response.

#### Noise reduction as a function of scan and spatial averaging

As expected, averaging over scans reduces the magnitude of the noise in the time series. For each subject, we computed the observed noise level of the time series (the standard deviation) averaged across all voxels in V1-V3 in a single scan, then computed the same metric on time series averaged across 8-10 scans (depending on subject). We found that the average noise level of all subjects decreased from 0.55% signal change (+/- 0.06%, std across 5 subjects) in a single scan to 0.19% (+/- 0.05%) after averaging the scans together.

The drop in noise magnitude for individual subjects as a function of scans averaged (gray curves, Fig 10a) appears close to a square-root-of-n decrease (dashed black curve, adjusted r-squared = 0.82). To confirm this drop-off, for each subject, we computed a ratio of the observed noise level when averaged over all scans to the expected noise level if all scans were independent (and thus would exhibit a square-root-of-n decrease in magnitude). When averaging across scans, the mean of the distribution of ratios across all subjects, evaluated at the maximum number of scans in each subject, was not significantly different from 1 (mean = 1.20, std = 0.24, p = 0.14, two-tailed t-test, n = 5 subjects), suggesting that noise between scans is independent. Hence, averaging across n-scans will decrease noise magnitude by a square-root-of-n.

**Figure 10:**
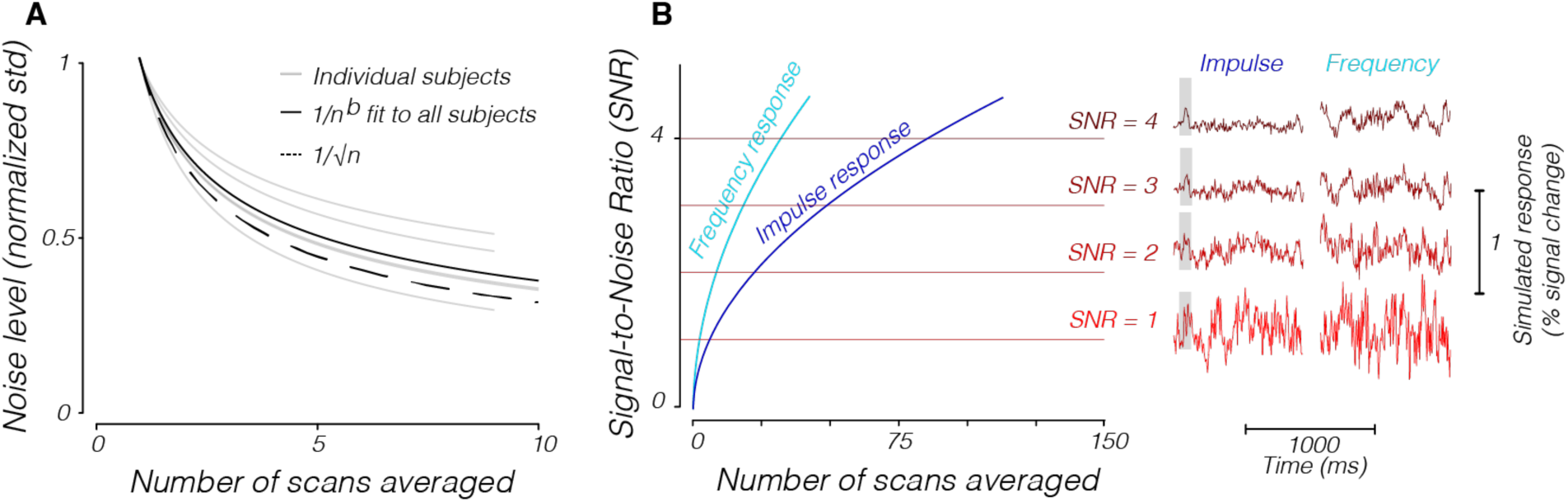
Noise magnitude and SNR estimates as a function of scan averaging. (A) The noise magnitude decreases as the number of averaged scans increases. The solid gray lines show the noise magnitude for individual subjects, and the solid black line shows the fit to all subjects. The dashed line shows the theoretical maximum noise reduction, which is the square root of the number of averaged scans. (B) The SNR increases as the number of averaged scans increases, as shown for both impulse (cyan) and frequency (dark blue, 5 Hz modulation) approaches. The right-hand side of the figure shows example time series for both approaches at certain SNRs. The time series are zero-meaned. In both cases, we assumed a single-scan noise level based on the average number of voxels in V1-V3 in our subjects (a standard deviation of 0.55% signal change from the mean).

Averaging over voxels in the human data also reduces the noise magnitude, but not by a square-root-of-n factor of the number of voxels included, suggesting that the variability in voxel time-series is spatially correlated. For each individual scan, we computed the standard deviation of the time series as a function of how many voxels we averaged over. Typical analyses examine contiguous voxels in particular regions of the brain and it is known that voxels are more correlated as a function of spatial proximity (Bergen and Jehee, 2018; Cohen and Kohn, 2011). Again, we calculated the ratio of the observed noise level for all voxels averaged to an ideal noise level that decreases as square-root-of-n with voxel inclusion. The mean of the distribution of these ratios across all scans is significantly above 1 (mean = 4.68, std. = 2.46, p = < 0.001, one-tailed t-test, n = 46 scans), indicating that there was correlated noise in the voxels.

#### SNR of a response to a briefly flashed stimulus (impulse response)

Using the relationships we calculated between the amount of scan- and spatial-averaging and the magnitude of the noise, we calculated the number of scans one would need to average over to achieve specific SNR levels when trying to measure a single-peaked response to a brief stimulus presentation (an impulse response). SNR was calculated as the magnitude of the peak of the response divided by the standard deviation of the noise time series. We estimated the strength of the stimulus response as 0.2% at its peak, which is close to the strongest responses reported in Toi et al. (Fig 1d, Toi et al.). To make noise estimates, we took the average noise magnitude (the standard deviation of the time series) for a single scan, averaged across all voxels in V1-V3 in our data. We then scaled that single-scan noise level as a square-root-of-n of the number of scans included (for details, see methods: SNR estimates). Comparing this scaled noise level as a function of scans to the expected response magnitude gives us expected SNR as a function of scans averaged (Fig 10b).

While response magnitude and noise will vary depending on many factors, including species differences and field strengths, this plot can easily be reinterpreted to accommodate such differences. For instance, if the response magnitude is expected to be twice as large, or the noise is half of what we report here, the SNR estimates would simply need to be multiplied by a factor of 2.

#### SNR of a response to a temporally modulating stimulus (frequency response)

Sinusoidally modulated stimulus designs can be constructed to improve SNR by concentrating the responses to specific frequencies outside of those in which known artifacts occur (Boynton et al., 2011; Engel et al., 1997; Norcia et al., 2015; Regan and Regan, 1988). Because the noise we measured had low amplitude in higher frequencies, a stimulus placed in those high frequencies would be more distinguishable from the noise. We calculated the number of scans one would need to average over to achieve specific SNR levels when trying to measure a response to a stimulus that is presented at a specific frequency, akin to the calculations we made for the impulse response case above (Fig 10b).

We simulated a 5 Hz response (0.2% signal change, peak to trough, as in the impulse response case) on top of 1/f noise and found that it was easier to detect than an impulse response under the same noise assumptions. The 5 Hz response frequency was chosen to be outside of the range of noise based on the 1/f drop-off, but is well within the range of temporal frequencies at which robust neural responses can be measured (Foster et al., 1985). SNR was defined as the amplitude of the response frequency divided by the noise level at that frequency, which was approximated as the average magnitude of the adjacent components (see methods: SNR estimates). With the same assumptions of signal amplitude and noise estimates taken from our data, a sinusoidally modulated response in time leads to better SNR than trying to detect a single-peaked response (Fig 10b).

As in the impulse case, the SNR curve scales relative to the assumptions made about signal and noise magnitude. In both cases, doubling the expected response to a stimulus would scale SNR curves by a factor of two, as would halving the amount of expected noise. Thus, regardless of the specific assumptions about the response and noise magnitude, because of the 1/f noise properties it will always be easier to achieve high SNR measuring a frequency response than measuring a single peak.

## Discussion

The conceit of the line-scan method is that it provides extremely high temporal resolution, even though it takes several orders of magnitude longer than EPI to acquire a full image. In a living animal, it is likely that there will be variations in the imaged substrate during the time required to collect an image. We show via simulations that these variations cause artifacts that are spread along the phase-encoding direction of the reconstructed images. These simulated artifacts qualitatively match data collected from human subjects. The most salient point of agreement between the simulation and data is the presence of structured temporal modulation in voxels outside of the brain and skull along the phase-encoding dimension. Another point is the smooth variation in the timing of the artifacts across the reconstructed images (phase). This agreement between the data and simulations suggests that the artifacts we observed in our data were caused by nonstationarities in the brain and skull between different line acquisitions over the course of each scan.

### Potential physiological sources of noisy modulations in humans

We did not see similar artifacts in the phantom, whose MR property is constant over time. This suggests that temporal modulations in the data arise from modulations of the substrate due to biological sources, including respiration, head motion, and the flow of blood and cerebrospinal fluid. The data are consistent with substrate modulations that vary smoothly across space, as evidenced from the slow changes in the phase of the first frequency component across the phase-encoding direction of the reconstructed images. The simulation supports this idea: degrading the spatial coherence of the substrate modulation (Fig 9b) changes the spatial pattern of the phase modulations, rendering them more random.

Eliminating the spreading of noisy modulations across the image would be near impossible. Take, for example, the cardiac cycle, which induces substantial variation in the MRI signal level (Chang et al., 2009; Chen et al., 2020; Hermes et al., 2023; Shmueli et al., 2007). We found that cardiac gating did not eliminate the spreading artifacts (Fig 6). While there may be some improvement that we were unable to detect with a small sample size, clearly the spreading effects remained even when gating to the cardiac cycle. It is possible that improvements to the gating procedure may improve its efficacy. For example, due to variability between heartbeats, we may have been unable to achieve perfect physiological gating which in theory would eliminate the spread of modulations from the heartbeat. However, gating to one physiological factor means you cannot gate to others, and it is likely that the modulations arise from multiple sources, practically ensuring spreading of the other modulations the experimenter did not gate to. In this sense, some amount of spreading is inevitable.

Our data suggest that to optimize the ability of line-scanning techniques to measure signals, DIANA or otherwise, underlying noisy modulations in the substrate should be minimized. Even if perfect gating to physiological cycles was achievable, gating would not eliminate the underlying temporal modulation in the brain. Rather, it would concentrate the modulations inside the brain instead of across the image.

Because the long line-scan acquisition time makes these physiological modulations more problematic than in EPI, and because these modulations cannot be fully addressed with conventional gating, humans may be a particularly difficult species in which to measure a DIANA response. The technical limitations of controlling physiological modulations in human subjects mean that the noise we have documented here will be hard to eliminate. In certain animal preparations, however, the experimenter may have greater control over physiological modulations and take actions to minimize them. These noisy modulations can be minimized during data collection by maintaining tight control over physiological state with anesthetic agents, respiratory cycles with artificial ventilation, and head motion with stereotaxic setups.

### Measuring a response

The difficulty in controlling these noisy modulations suggests that the best approach to picking up an electrophysiological signal is to use a stimulus that evokes a response that is unlike the noise, rather than trying to eliminate the noise altogether. This can be done by targeting temporal frequencies with minimal noise and maximal response. The 1/f noise structure suggests that increasing the targeted response frequency will decrease the noise. The selection of a response frequency, however, must also account for the fact that cortical neurons are better tuned to lower temporal frequencies. We found that a 5 Hz stimulus is high enough frequency to be outside of the major noise terms, and is still in the 4-10 Hz range that generates a large neural response in early visual cortex (Foster et al., 1985; Hawken et al., 1996; Horiguchi et al., 2009; Stigliani et al., 2017; Sun et al., 2007).

### Contrast adaptation and SNR

While neural responses to a repeated stimulus decline due to effects such as repetition suppression (Dobbins et al., 2004; Grill-Spector et al., 2006) or contrast adaptation (Gardner et al., 2005; Kohn and Movshon, 2003; Ohzawa et al., 1985, 1982), our analysis and modeling suggest that the typical strategy of averaging over repeats can overcome signal loss due to adaptation. In the various attempts to measure the DIANA signal, stimuli have typically been presented very briefly (tens of ms) with long intervals (hundreds of ms) in between each presentation (Choi et al., 2023; Toi et al., 2022). Adaptation in sensory visual neurons can occur at multiple timescales from milliseconds (Lisberger and Movshon, 1999) to seconds (Bonds, 1991) to days (Bao and Engel, 2012; Engel, 2019) and recovery from adaptation typically occurs with time constants that are in the same order of magnitude as the rate of adaptation (for review, see (Kohn, 2007)). That is, given the order of magnitude difference between the length of sensory stimulation and the recovery time between presentations typically used for line-scanning, responses would be expected to be nearly, if not fully, recovered by the next line and therefore produce negligible artifacts.

Additionally, for adaptation to result in loss of signal with more averaging, adaptation would have to decrease the signal faster than the noise decreases with averaging. We confirmed that the standard deviation of the noise in our scans decreased proportional to the square root of the number of scans as is expected for independent and identically distributed noise. As explained above, neural adaptation for the stimulus paradigms typically used with line-scanning techniques is not expected to decrease the signal faster than the square root of the number of scans, and thus signal-to-noise ratios would only be expected to increase with more averaging.

### Potential neural sources of nonstationarity

Contrast adaptation is a nonstationarity in the response across multiple stimulus presentations and thus between line acquisitions. Like the nonstationarities analyzed here, contrast adaptation will spread the response in the phase-encoding direction. In human visual cortex, fMRI measurements show that the response to repeated stimuli decreases gradually by about 20-30% (Grill-Spector, 1999; Murray et al. 2002) in a localized region. Thus, we do not expect a substantial impediment to measuring neural signals with line-scanning sequences.

As adaptation is a gradual process, often modeled as an exponential drop-off in response magnitude, the difference in signal magnitude between adjacent line acquisitions is small. These gradual changes are especially small compared to the changes we modeled in this paper where we abruptly change the phase of the underlying signal. Unpublished simulations show that the amount of spreading in the reconstructed image is a function of the magnitude of the difference of the imaging substrate between adjacent line acquisitions. Thus, while we expect that contrast adaptation will spread a neural response across the image, that spreading will be minimal because of the slow dynamics of the adaptation.

While we expect spreading due to adaptation (or variability in response onset) to be small for reasons described above, it may still be the case that these nonstationarities cause responses from one position in the brain to manifest as responses in different locations, perhaps even shifted in time. This possibility is similar to the emergence of systematic timing differences that we observe via the phase gradients in the sinusoidal modulation simulations, in which a signal that modulates the entire brain at once appears shifted in time across voxels that did not initially modulate in the reconstruction. It is often of interest to examine responses across space and time, for example when examining signal propagation from thalamus to cortex (as in Toi et al.) or in the cortical hierarchy. Groups attempting to show such timing differences between different regions of the brain should ensure that they are positioned orthogonal to the phase-encoding direction of the image acquisition to ensure that timing effects are not confused with signal spreading in the reconstruction.

## Conclusion

Direct measurement of spatially localized and temporally resolved neural activity in the living human brain would enable many new discoveries. This possibility justifies the excitement, and scrutiny, evoked by Toi et al. The failures to replicate the original findings (Choi et al., 2023; Hodono et al., 2023; Thorp, 2023) have been met by a defense from the original authors. Our efforts to find a signal in humans at 3T did not succeed, prompting an analysis of the noise characteristics that one would expect using the original line scan methods.

Several of the basic features of the noise were expected given the long history using the line-scan method for cardiac imaging. The data suggest that our measurements were made in the presence of global modulations in the brain and skull which are asynchronous between line acquisitions. This spreads the low-frequency-weighted temporal modulations across the image. The temporal characteristics of these noisy modulations make them easily confusable with certain classes of stimulus responses, including the signal one might expect from a briefly presented stimulus. Both the measurements and a simulation of the line-scan acquisition of a globally modulating brain substrate confirmed this expectation.

Our results suggest that noisy physiological modulations are particularly problematic in line-scan MRI because of the long acquisition times - the lines in any single image are collected over roughly a minute. The line-scan spreading artifacts we encounter are substantial but easy to diagnose. A metric like the autocorrelation permutation diagnoses the spreading, both of signals and noise. Further, the spreading can be diagnosed from the regular shift over space of the phase of the modulation frequency. Modifications to k-space ordering and trajectories, combined with novel forms of navigators and field monitoring may help suppress the artifacts we’ve shown here. However, in light of the difficulties in minimizing the noise, we suggest alternative approaches to increase one’s ability to detect a response. We show that averaging is an effective way to reduce noise, and that alternative stimulus paradigms would increase SNR in the presence of the noise. While we were unable to detect a neural response to a brief stimulus in humans with a 3T scanner, more controlled animal preparations with higher field strengths, more efficient stimuli, and more averaging may have more success.

## Acknowledgements

We thank Brian Rutt, Kawin Setsompop, Michael Zeineh, and Eli Merriam for useful discussions and their insights into the methods. We are grateful to Austin Kuo, Akshay Jagadeesh, Jiwon Yeon, Hyunwoo Gu, Josh Ryu, and Alex Durango for their help with data acquisition.

## Contributions

All authors contributed to the designed experiments. J.W. and H.W. collected the data. J.W. created the simulations. J.W., B.W., and J.L.G. analyzed the data and wrote the manuscript.

## Data and code availability

Data and analyses are available on osf at https://osf.io/243pn/. Simulation and analysis code is available at https://github.com/joshmw/emri. Some analysis code requires scripts from https://github.com/justingardner/mrTools and https://github.com/justingardner/gru.

